# Chromosome-scale genome assembly of *Apocynum pictum*, a drought-tolerant medicinal plant from the Tarim Basin

**DOI:** 10.1101/2024.03.08.584125

**Authors:** Wenlong Xie, Baowei Bai, Yanqing Wang

## Abstract

*Apocynum pictum* Schrenk is a semi-shrub of the Apocynaceae family with a wide distribution throughout the Tarim Basin that holds significant ecological, medicinal, and economic values. Here, we report the assembly of its chromosome-level reference genome using Nanopore long-read, Illumina HiSeq paired-end, and high-throughput chromosome conformation capture sequencing. The final assembly is 225.32 Mb in length with a scaffold N50 of 19.64 Mb. It contains 23,147 protein-coding genes across 11 chromosomes, 21,148 of which (91.36%) have protein functional annotations. Comparative genomics analysis revealed that *A. pictum* diverged from the closely related species *Apocynum venetum* approximately 2.2 million years ago and has not undergone additional polyploidizations after the core eudicot WGT-γ event. Karyotype evolution analysis was used to characterize interchromosomal rearrangements in representative Apocynaceae species and revealed that several *A. pictum* chromosomes were derived entirely from single chromosomes of the ancestral eudicot karyotype. Finally, we identified 50 members of the well-known stress-responsive WRKY transcription factor family and used transcriptomic data to document changes in their expression at two stages of drought stress, identifying a number of promising candidate genes. Overall, this study provides high-quality genomic resources for evolutionary and comparative genomics of the Apocynaceae, as well as initial molecular insights into the drought adaptation of this valuable desert plant.

## Introduction

The Tarim Basin, situated in southern Xinjiang, is the largest inland basin in China. It is bordered by the Kunlun Mountains in the south, connected to the Tianshan Mountains in the north, and stretches from the Altun Mountains in the east to the Pamir Plateau in the west. In the central part of the basin lies the Taklamakan Desert. The Tarim Basin falls within the temperate zone and experiences a continental arid and semi-arid climate. It frequently encounters dust storms and sandstorms and is characterized by long periods of sunshine and significant temperature fluctuations between day and night. Previous studies have concluded that a hyperarid climate has persisted within the basin for at least 5.3 million years, and there is evidence of increasing aridity^1^. The basin’s unique geographic location and challenging climate conditions contribute to a mere 10% vegetation coverage along its periphery, primarily composed of shrubs and semi-shrubs^1,2^. These vegetation types are excellent materials for studying plant adaptations to dry environments.

*Apocynum pictum* Schrenk (2n = 22, synonym of *Apocynum hendersonii* Hook. f.), commonly referred to as Bai ma, is a semi-shrub from the Apocynaceae family that grows predominantly in the northwestern inland region in areas surrounding saline-alkali wastelands, desert edges, and river alluvial plains. It exhibits a wide distribution throughout the Tarim Basin^3^. *A. pictum* shows strong ecological adaptability to cold, drought, wind erosion, and saline-alkaline conditions, and it is often used as a windbreak and sand-fixing plant in Xinjiang, China. *A. pictum* also serves as a valuable source of honey and has economic significance as an excellent fiber plant because of the high quality of phloem fibers in its stems^4^. In addition, *A. pictum* has been used as a pharmacological plant because of its significant content of flavonoids, primarily quercetin^5^. Previous studies have focused on the closely related species *Apocynum venetum*, also called Luobuma, which has demonstrated a wide range of medicinal properties^6^. Although recent developments in sequencing technology have enabled assembly of the chloroplast and nuclear genomes of *A. pictum*^7,8^, there is still a need for a high quality, chromosome-scale reference genome.

In this study, we produced a chromosome-scale reference genome and annotations for *A. pictum* by integrating Oxford Nanopore Technologies (ONT) long-read, Illumina HiSeq paired-end, and high-throughput chromosome conformation capture (Hi-C) sequencing. We examined the general characteristics of *A. pictum* genome evolution, identified 50 members of the WRKY transcription factor family, and performed transcriptome sequencing at two stages of the drought stress response. The *de novo* assembly and annotation of this high-quality *A. pictum* genome provide a basis for future evolutionary research on the Apocynaceae family and support continued study and utilization of this versatile and economically valuable plant.

## Results

### Genome survey, sequencing, assembly, and assessment

We estimated the genome size of *A. pictum* by analyzing the distribution of 17-mers using 16.98 Gb of Illumina HiSeq data. The resulting plot showed three distinct peaks: a primary peak with a depth of 90, and a secondary peak and smaller peak with half and double the depth of the primary peak, respectively, indicative of a heterozygous peak and a homozygous peak (Figure 1B). The *A. pictum* genome was estimated to be ∼202 Mb in size, with a heterozygosity rate of 1.13% and 37% repetitive sequences.

**Figure 1.**
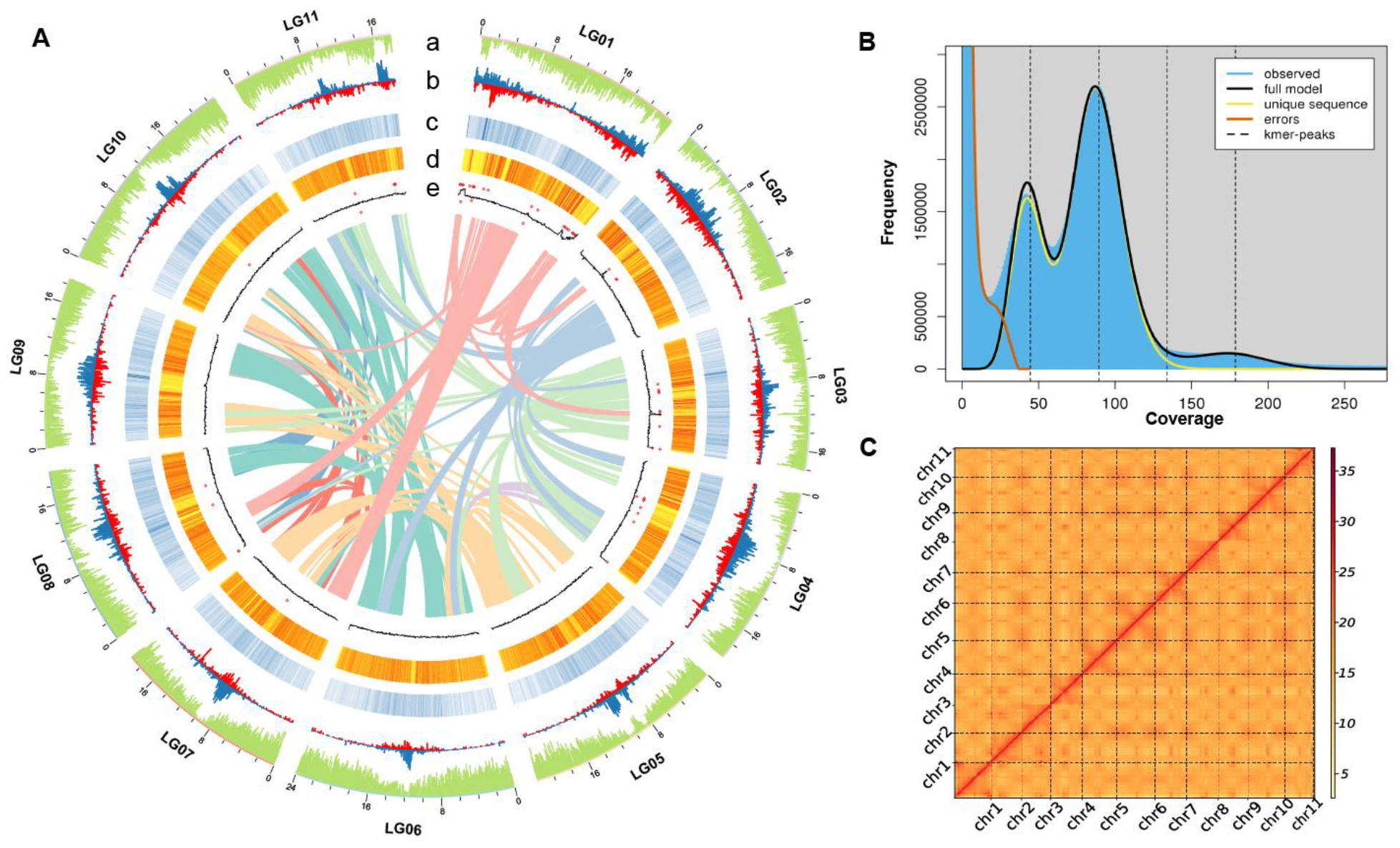
(A) Landscape of the *A. pictum* genome assembly and annotation. Tracks from outside to inside correspond to a, gene density; b, Copia density in blue and Gypsy density in red; c, repeat density; d, SNP density; e, GC density (black line) and N ratio (red symbols). Genome features are depicted in 10-kb windows across the chromosomes. (B) 17-mer depth distribution. The genome was estimated to be 202 Mb in size, with a heterozygosity rate of 1.13% and a repeat percentage of 37%. (C) Hi-C assisted assembly of *A. pictum* pseudochromosomes. The heatmap reveals the presence of an anti-diagonal pattern of intrachromosomal interactions, which were scaffolded and assembled independently.

We used a combination of ONT, Illumina HiSeq, and Hi-C data to assemble the *A. pictum* genome (Supplementary Table S1). We used 21.49 Gb of ONT long reads (95× coverage) for self-correction and initial contig assembly, then polished the resulting assembly twice using Illumina short reads. This produced a preliminary assembly of 59 contigs with a contig N50 of 9.63 Mb (Table 1). We used 48.26 Gb of Hi-C data to obtain 26 scaffolds with a scaffold N50 of 19.64 Mb, which enabled us to anchor the preliminary contigs to eleven pseudochromosomes (Figure 1A, C). Nearly all (99.55%) of the base pairs were successfully anchored to pseudochromosomes, producing a final assembled genome of 225 Mb, slightly larger than the size predicted by the genome survey. This discrepancy may be attributed to the high heterozygosity and repeat content of the genome.

**Table 1.** Statistics for *A. pictum* genome assembly and annotation.

| Feature | Value |
| --- | --- |
| Genome size (bp) | 225,320,346 |
| GC content (%) | 32.44 |
| Number of contigs | 59 |
| Contig N50 (bp) | 9,635,369 |
| Contig N90 (bp) | 3,354,232 |
| Number of scaffolds | 26 |
| Scaffold N50 (bp) | 19,644,087 |
| Scaffold N90 (bp) | 18,080,797 |
| Sequence anchored to pseudochromosomes (bp) | 224,312,087 |
| LAI | 15.37 |
| Repetitive sequences (%) | 35.95 |
| Gene number | 23,147 |
| Average gene length (bp) | 3,574 |

The completeness of the assembled genome was assessed using Benchmarking Universal Single Copy Orthologs (BUSCO)^9^. Of the 2,326 BUSCO orthologs in the eudicots database, 2,250 (96.8%) were completely captured in the genome (Supplementary Table S2). To assess genome accuracy, we mapped clean Illumina short-read data to the genome, and 93.41% of the reads were properly paired. We then identified 9,033 homozygous single nucleotide polymorphisms (SNPs) and 15,965 homozygous insertion– deletions (indels) based on the mapping results. The assembly accuracy rate based on the percentage of homozygous single-nucleotide variants was 99.99% (Supplementary Table S3). Assembly quality was also assessed by calculating the Long Terminal Repeat (LTR) Assembly Index (LAI). The LAI value of 15.37 fell between 10 and 20, indicating that the *A. pictum* genome was of reference quality^10^.

### Repetitive element and gene annotations

A total of 35.95% (80.99 Mb) of the assembly was identified as repetitive elements using both *de novo* and homology-based methods (Table 1). This included tandem repeats, which represented 2.64% (5.94 Mb) of the total genome (Supplementary Table S4). Another category comprised elements that were dispersed across the genome, primarily transposable elements (TEs), which represented 22.35% (50.36 Mb) of the genome. Among the LTR retrotransposons (LTR-RTs) of the class-I TEs, *Gypsy* (9.96%) and *Copia* (8.69%) were the most prevalent super-families (Supplementary Table S5). We also identified 1,827 non-coding RNAs in the *A. pictum* genome, including 85 miRNAs, 549 snRNAs, 416 tRNAs, and 688 rRNAs (Supplementary Table S6).

We used transcriptome-based, homology-based, and *de novo* strategies to predict protein-coding genes in the assembled genome. We identified 23,147 protein-coding genes mapped to the 11 pseudochromosomes with an average length of 3,574 bp and an average CDS length of 1,303 bp (Supplementary Table S7). Most (21,148; 91.36%) of the predicted protein-coding genes were functionally annotated using the EggNOG^11^ database, and 81.38% contained conserved protein domains as determined using the PFAM^12^ database. These results provide valuable information for gene family identification (Supplementary Table S8). We assessed the quality of the protein-coding gene annotations using BUSCO and obtained 2,232 (96.0%) of the complete BUSCO orthologs (Supplementary Table S2).

### Phylogenetic analysis

We performed maximum likelihood phylogenetic analyses using protein sequences from 10 species: two from Apocynoideae (*Apocynum pictum* and *Apocynum venetum*), two from Rauvolfioideae (*Catharanthus roseus*^13^ and *Voacanga thouarsii*^14^), three from Asclepiadoideae (*Marsdenia tenacissima*^15^, *Asclepias syriaca*^16^, and *Calotropis gigantea*^17^), two from the Rosids (*Arabidopsis thaliana* and *Vitis vinifera*), and one from the monocots (*Oryza sativa*) as an outgroup species (Supplementary Table S9). Apocynoideae, Rauvolfioideae, and Asclepiadoideae are subfamilies of the flowering plant family Apocynaceae, and the rosids *A. thaliana* and *V. vinifera* were used to calibrate the divergence times of the species tree. On the basis of protein sequence homology, 244,695 (91.3%) genes were clustered into 24,459 gene families; 13,138 families were shared by subfamilies of the Apocynaceae, and 1,077 families were specific to the Asclepiadoideae (Figure 2C,D). *A. pictum* and *A. venetum* differ from the other five Apocynaceae species in being widely distributed in salt-barren zones and desert steppes^3^, suggesting that their unique gene families may include genes associated with stress tolerance.

**Figure 2.**
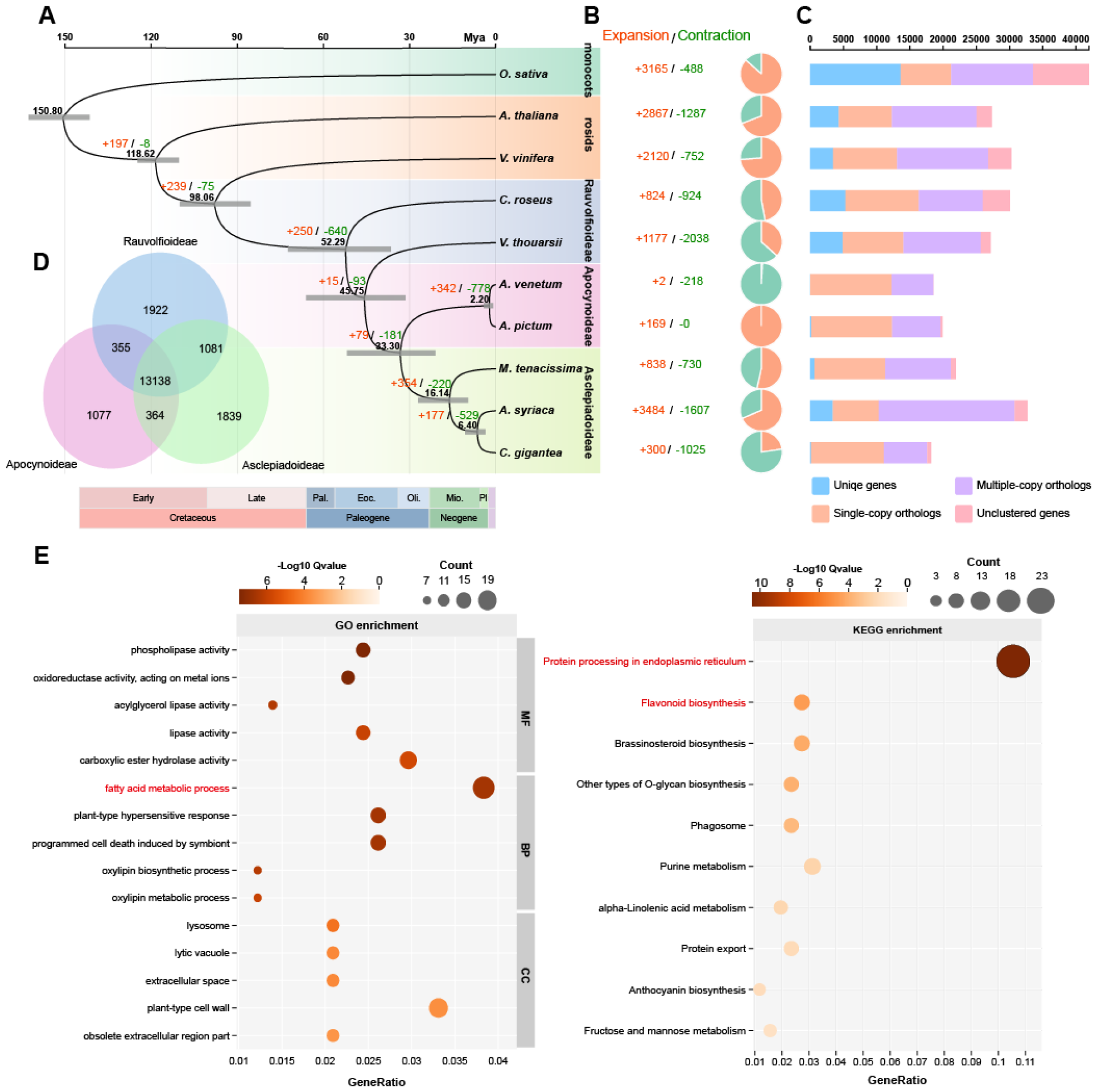
(A) Maximum likelihood phylogenetic tree constructed from 1,376 single-copy orthologs and divergence-time estimates for *A. pictum* and other species, rooted with *O. sativa* as the outgroup. (B) Expansions and contractions of gene families in each species. (C) Numbers of unique, single-copy, and multi-copy genes in *A. pictum* and other species. (D) Venn diagram of unique and shared orthologous genes among Rauvolfioideae, Apocynoideae, and Asclepiadoideae. (E) GO and KEGG enrichment analysis of expanded gene families in *A. pictum*, related to the common ancestor of the *A. pictum* and *A. veneum*. The top five significantly enriched biological process (BP), cellular component (CC), and molecular function (MF) GO terms are shown at left and the top 10 significantly enriched KEGG pathways at right. Dot color represents the Q-value, and dot size represents the number of genes mapped to the indicated terms/pathways.

The family Apocynaceae has been classified into five subfamilies^18^. To date, high-throughput genome sequencing has been performed on species from three of these subfamilies. Here, we examined the evolutionary patterns of representative Apocynaceae species based on a phylogenetic tree derived from 1,376 single-copy orthologs (Figure 2A). The findings indicate that Rauvolfioideae is the closest relative to the ancestral lineage, suggesting its early formation within the evolutionary lineage of Apocynaceae species. Asclepiadoideae appeared to have diverged from a hypothetical common ancestor approximately 33.3 million years ago (Mya). This divergence time aligns with the fossil evidence, in which the earliest record of Asclepiadoideae dates back to the Eocene epoch (33.9–56 Mya)^19^. Asclepiadoideae and Apocynoideae are part of the APSA clade (Apocynoideae, Periplocoideae, Secamonoideae, and Asclepiadoideae); we estimated the age of the crown node for the APSA at 45.75 Mya, consistent with estimates from previous reports (45–65 Mya)^18,20^.

*A. pictum* and *A. venetum* display close genetic relationships and were estimated to have diverged from a common ancestor 2.2 Mya, after the divergence of Asclepiadoideae at 33.3 Mya. The global cooling and aridification that began at the Eocene–Oligocene climate transition (34 Mya) led to the expansion of temperate regions and the establishment of modern grasslands and deserts^21^. We speculate that these changes may have facilitated the establishment and subsequent expansion of the *A. pictum* and *A. venetum* ancestor. In comparison to their ancestor, *A. pictum* showed an expansion of 169 gene families, and *A. venetum* exhibited a contraction of 218 gene families and expansion of 2 gene families (Figure 2B). We examined the functions of the expanded gene families in *A. pictum* by Gene Ontology (GO) and Kyoto Encyclopedia of Genes and Genomes (KEGG) enrichment analysis (Figure 2E). The expanded genes were most highly enriched in the GO term “fatty acid metabolic process” (GO:0006631) and the KEGG pathway “protein processing in endoplasmic reticulum” (associated with 27 genes), which may be related to environmental stress responses in plants^22,23^. The second most significantly enriched pathway was “flavonoid biosynthesis,” which could account for the differences in overall content and composition of flavonoids observed between the two species^24^.

### WGD analyses and karyotype evolution

We used synonymous substitution rates (Ks) to investigate the polyploidization histories of *A. pictum* and two other Apocynaceae species with high-quality chromosome-scale genome assemblies, *M. tenacissima*^15^ from the Asclepiadoideae and *C. roseus*^13^ from the Rauvolfioideae (Figure 3A). We estimated the Ks distribution of orthologous gene pairs in intergenomic syntenic blocks and observed distinct narrow peaks at 0.36 and 0.46, suggesting that *M. tenacissima* diverged from the common ancestor after the divergence of *A. pictum* and *C. roseus*. In genomic synteny plots of the three species (Figure 3B), the individual chromosomes of *A. pictum* exhibited the most significant matches with corresponding chromosomes of *M. tenacissima*, suggesting a closer genetic relationship of *A. pictum* with *M. tenacissima* than with *C. roseus*. When we estimated the Ks distribution of paralogous gene pairs in intragenomic syntenic blocks, we detected a single broad peak at approximately 1.95, indicating that a single polyploidization event was shared by all three species.

**Figure 3.**
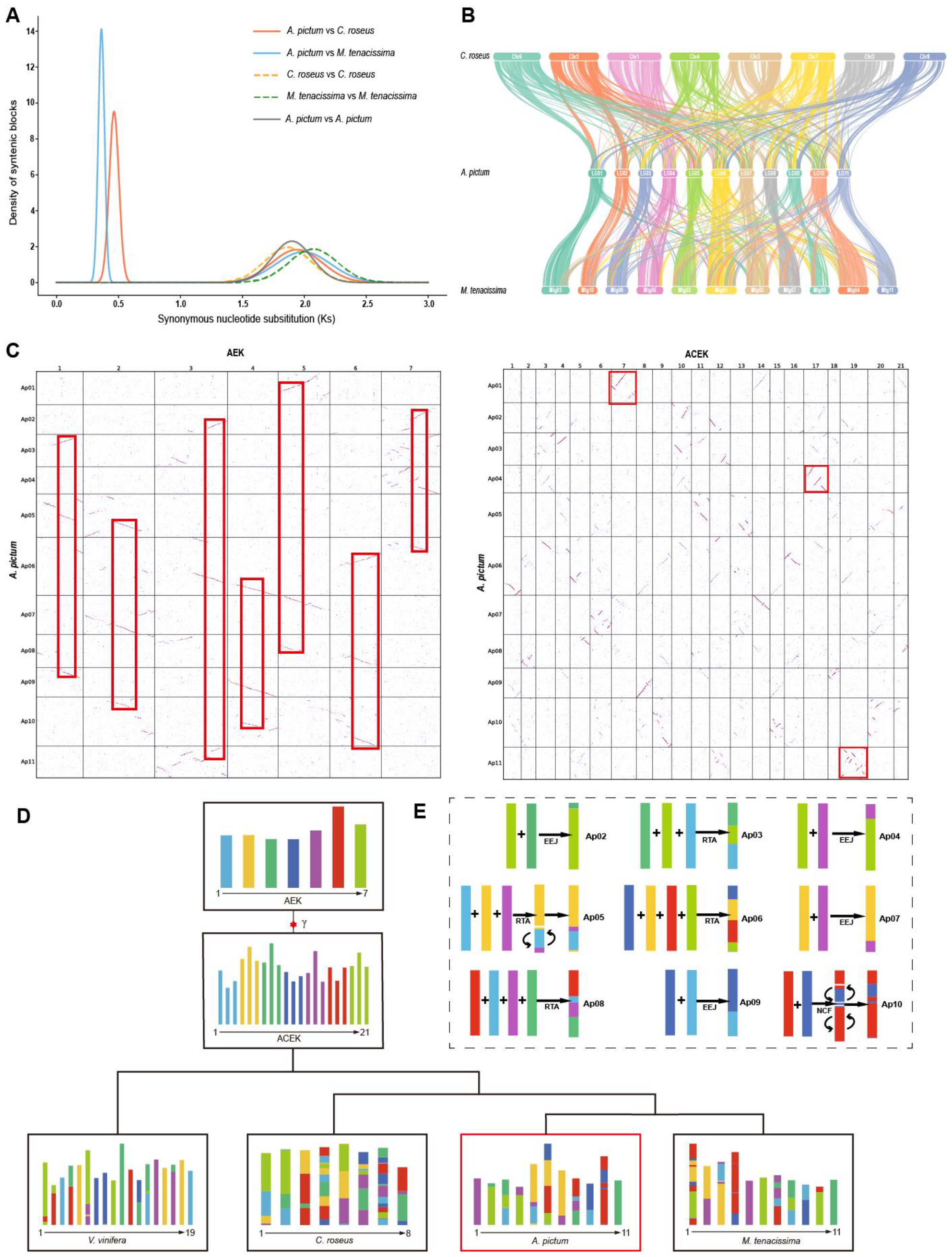
(A) Synonymous nucleotide substitution (Ks) distributions for different pairs of *A. pictum, C. roseus*, and *M. tenacissima*. The narrow blue peak represents the *A. pictum* and *M. tenacissima* divergence at Ks = 0.36, and the narrow orange peak represents the *A. pictum* and *C. roseus* divergence at Ks = 0.46. The broad peak around Ks = 1.95 represents the shared WGT-γ event in the three species. (B) Gene synteny between *A. pictum, C. roseus*, and *M. tenacissima*. (C) Syntenic block dotplots between *A. pictum* and the AEK (depth ratio 3:1) and between *A. pictum* and the ACEK (depth ratio 1:1). Red lines represent high-confidence syntenic blocks. Chromosomes 1, 4, and 11 of *A. pictum* are almost entirely derived from chromosomes 7, 17, and 19 of the ancestral ACEK. (D) Karyotype projections of *V. vinifera, C. roseus, A. pictum*, and *M. tenacissima*. The AEK gave rise to the ACEK through the WGT-γ event. (E) Proposed history of karyotype evolution in *A. pictum*. RTA, reciprocally translocated chromosome arms; EEJ, end-to-end joining; NCF, nested chromosome fusion.

To characterize the whole-genome duplication (WGD) event in the evolutionary history of *A. pictum*, we performed karyotype assessments using the ancestral eudicot karyotype (AEK) and the ancestral core eudicot karyotype (ACEK) as references^25^. Previous studies have revealed that the AEK, which consists of 7 protochromosomes, experienced a whole genome triplication event (WGT-γ) that resulted in formation of the ACEK, with 21 chromosomes^26^. We mapped the *A. pictum* genome onto the AEK and ACEK genomes and generated syntenic block dotplots (Figure 3C). It was evident that each protochromosome in the AEK had three highly homologous regions in the *A. pictum* genome, indicating a syntenic depth ratio of 3:1. By contrast, the syntenic depth ratio between the ACEK and *A. pictum* was 1:1, confirming that *A. pictum* experienced the eudicot WGT-γ and did not undergo any additional polyploidization events, similar to the model plant *V. vinifera*^27^. Our analyses showed that chromosomes 1, 4, and 11 of *A. pictum* originated from ancestral chromosomes 7, 17, and 19 of the ACEK (Figure 3C). This finding implies that there has been limited rearrangement of these chromosomes since the WGT-γ event, suggesting that certain genes present in these conserved chromosomes may play roles in development and may have prevented chromosomal fusion.

We next performed a karyotype evolution analysis to examine details of chromosome reorganization in representative Apocynaceae. We identified 335, 368, and 354 homologous blocks between *A. pictum* and the AEK, *C. roseus* and the AEK, and *M. tenacissima* and the AEK, respectively. In Figure 3D, chromosomal regions of extant species that are homologous to portions of the AEK chromosomes are indicated with corresponding colors (without considering any gene loss). It is clear that *C. roseus* has undergone more chromosome fusions than *A. pictum* and *M. tenacissima*. For instance, every chromosome in *C. roseus* originated from multiple AEK ancestral chromosomes, whereas chromosomes 1 and 11 in *A. pictum* and chromosomes 5, 6, and 11 in *M. tenacissima* originated from single AEK ancestral chromosomes (Figure 3D). Fusion is the predominant form of interchromosomal rearrangement, encompassing reciprocally translocated chromosome arms (RTAs), end-to-end joining (EEJ), and nested chromosome fusion (NCF). The dotplot and karyotype projection of *A. pictum* and the AEK reveal that chromosomes 2, 4, 7, and 9 in *A. pictum* were formed through EEJ from AEK protochromosomes; chromosomes 3, 5, 6, and 8 were formed through RTA; and chromosome 10 was formed through NCF (Fig. 3E). It is likely that chromosome 1 and 11 were retained as independent chromosomes inherited from the AEK.

### Identification of *A. pictum* WRKYs

The WRKY transcription factors (TFs), named after their conserved N-terminal WRKY motifs, play crucial roles in various physiological processes, particularly in responses to abiotic stresses like drought, temperature, ultraviolet radiation, and salinity^28^. Given their noted role in drought response, we identified and characterized members of this TF superfamily in the drought-tolerant shrub *A. pictum*. Using *Arabidopsis* WRKY sequences from the TAIR^29^ database as queries, we identified 50 putative WRKY genes in the *A. pictum* genome (Fig. 4A). The number of WRKY family members was lower in *A. pictum* than in *Arabidopsis* (over 70) but quite similar to that in the closely related species *Catharanthus roseus* (49)^13^. Expansion of the WRKY family in *Arabidopsis* may reflect additional whole-genome duplications in the Brassicaceae lineage (WGD-α and WGD-β). We named the *A. pictum* genes ApWRKY1 to ApWRKY50 on the basis of their order in the genome sequence from chromosome 2 to 11. We first performed a comprehensive bioinformatic analysis of their predicted proteins (Supplementary Table S10), which ranged from 167 to 737 amino acids (aa) in length, with molecular weights from 19,193 to 80,662 Da and isoelectric points (pIs) from 4.90 to 9.76. All were classified as unstable proteins because their instability indices were greater than 40; their aliphatic indices and GRAVY scores suggested a higher potential for hydrophilicity^30^. Analysis of collinearity among the WRKY genes revealed 15 segmentally duplicated pairs across the *A. pictum* genome (Figure 4B).

**Figure 4.**
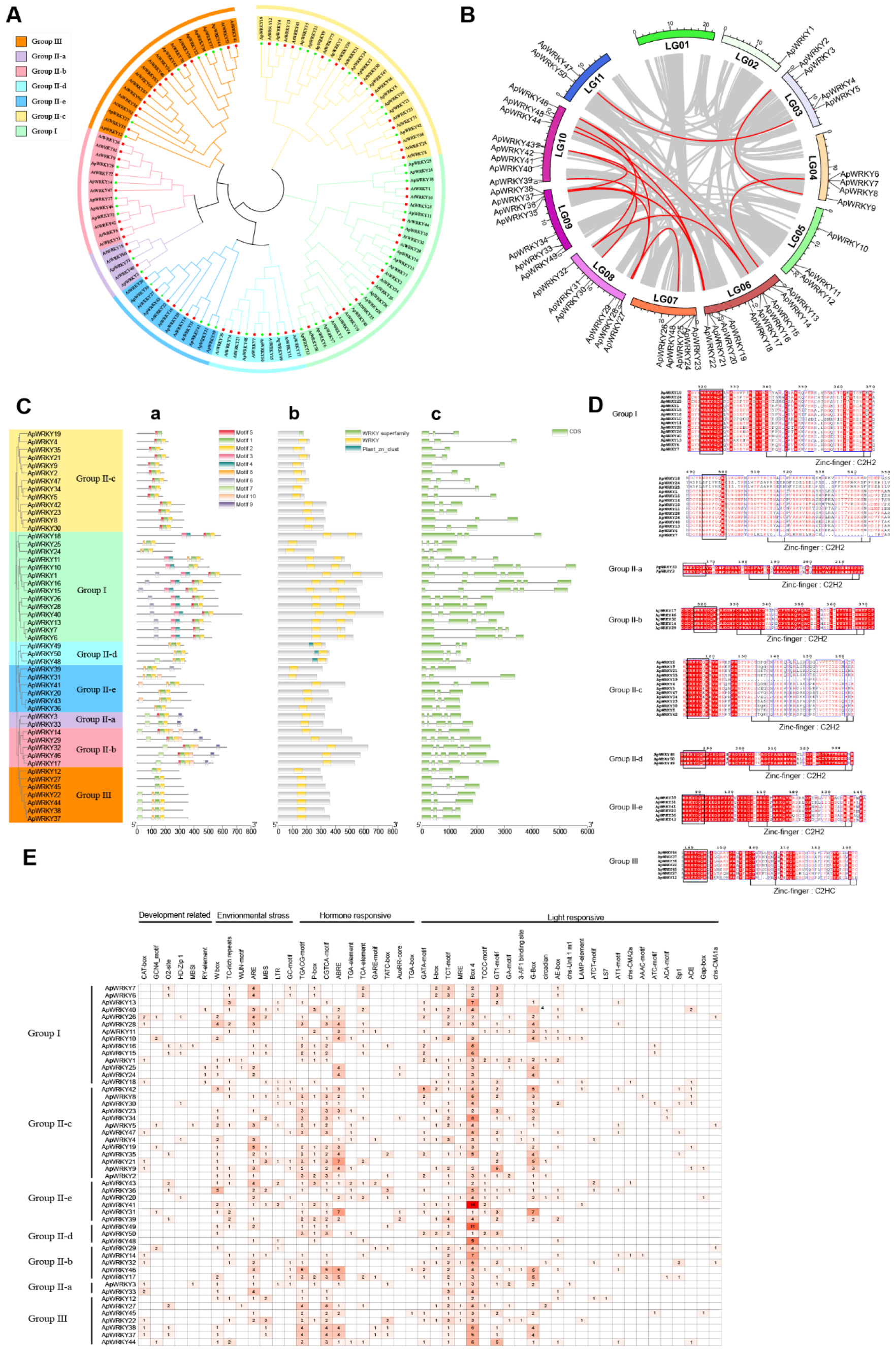
(A) Neighbor-joining phylogenetic tree of WRKY proteins from *A. thaliana* and *A. pictum*. The WRKY proteins were classified into three groups (I–III), and group II proteins were classified into five subgroups (IIa–IIe). (B) Distribution of *ApWRKY* genes on *A. pictum* chromosomes. Syntenic blocks are linked by gray lines, and syntenic *ApWRKY* gene pairs are linked with red lines. (C) Analysis of conserved protein motifs (a), conserved protein domains (b), and intron–exon structures of ApWRKY genes (c). Green boxes indicate exons. (D) The conserved WRKY domain consists of the WRKYQK sequence followed by a zinc finger structure. (E) Analysis of cis-acting elements in *ApWRKY* gene promoters. Cell color corresponds to the number of each element in the promoter of each *ApWRKY*.

Like the *Arabidopsis* WRKYs, the 50 ApWRKY proteins could be classified into three major groups^31^: group I (two WRKY domains and two C2H2 zinc-finger motifs), group II (one WRKY domain, one C2H2 zinc-finger motif), and group III (one WRKY domain, one C2HC zinc-finger motif) (Figure 4A). Group II could be divided into five subgroups (IIa–e), all of which contained *A. pictum* and *Arabidopsis* members.

Some of the ApWRKYs exhibited atypical variations in their conserved domains or zinc-finger structures. For example, although the WRKY DNA-binding domain typically consists of the heptapeptide WRKYGQK^32^, this sequence was WRKYGKK in ApWRKY5, 47, and 34. The typical group I zinc-finger motif (CX4C-HXH) was CX5C-HXH in ApWRKY24, 25, and 10, and the typical group III zinc-finger motif (CX7C-HXC) was CX4C-HXH in ApWRKY12 (Figure 4D). Several WRKYs appeared to have lost specific domains. In group I, ApWRKY24 and ApWRKY25 contained only one WRKY domain and one zinc-finger motif, and the N-terminal WRKY domain in ApWRKY7 had lost the zinc-finger motif. Similar loss of the zinc-finger structure was observed in ApWRKY19 from group II-c (Figure 4D). Previous work in rice showed that the WRKY family has been shaped by multiple episodes of domain acquisition and loss^33^, and some of the ApWRKY proteins identified here also appear to have experienced the loss of individual domains during evolution.

We next examined the conserved motifs and domains of the WRKY proteins and the intron–exon structures of the WRKY genes in the context of their phylogenetic relationships (Figure 4C). All ApWRKY proteins contained at least one of the two major DNA-binding domains, which were composed of motifs 1/2 and 3/4, respectively, in the MEME analysis (Supplementary Figure S1). In addition to the WRKY domain, the plant_zn_cluster domain was also found in ApWRKY48, 49, and 50 from group II-d. This domain is also present in group II-d WRKYs from other drought-resistant species such as *Caragana korshinskii* and *Caragana intermedia*^34,35^. The number of exons in ApWRKY genes ranged from 2 to 6, and genes from the same phylogenetic clade exhibited similar intron–exon structures (Figure 4C).

WRKY proteins bind to W-box elements in gene promoters to activate or inhibit transcription of downstream genes. They can also bind to their own promoters or to those of other *WRKYs* to create self-regulatory or cross-regulatory networks^36^. We therefore scanned the 2,000 bp upstream of each *ApWRKY* transcription initiation site to identify potential *cis*-acting elements. Forty eight elements were identified in the *WRKY* promoters (Figure 4E). Light-responsive elements were the most abundant. For example, the Box 4 element was present in almost all *ApWRKY* promoters except those of *ApWRKY11* and *21*. Elements related to five plant hormones were also identified: MeJA (TGACG-motif and CGTCA-motif), gibberellin (p-box, GARE-motif, and TATC-box), abscisic acid (ABRE), auxin (AuxRR-core, TGA-element, and TGA-box), and salicylic acid (TCA-element). *cis*-elements associated with stress-related hormones, such as the abscisic acid responsiveness element, ABRE, were particularly abundant in *ApWRKY* promoters. Seven stress-related *cis*-elements were also present, among which the W box, anaerobic induction element (ARE), and TC-rich repeats were particularly common. These results highlight the likely involvement of ApWRKYs in multiple physiological processes, particularly abiotic and biotic stress responses.

### Expression profiles of *ApWRKYs* under drought stress

Leaf tissues from *A. pictum* seedlings were subjected to PEG-induced drought stress for 0, 4, and 12 h, then sampled for transcriptome sequencing to investigate changes in *ApWRKY* expression under drought. Expression was quantified as transcripts per kilobase of exon model per million mapped reads (TPM), and the log2(TPM+1) values for each gene were hierarchically clustered and visualized in a heatmap (Supplementary Figure S2). We identified differentially expressed genes (DEGs) by comparing each drought time point (4 and 12 h) with the 0-h control.

We identified 3,225 DEGs, including 16 *ApWRKY* genes, that were classified into eight clusters (C1–C8) on the basis of their temporal expression patterns (Figure 5A,C). Five *ApWRKYs* (*ApWRKY44, 24, 33, 22*, and *3*) were present in C5, whose 481 members continued to increase in expression as the duration of drought stress increased. GO and KEGG enrichment analyses revealed that DEGs in this cluster were significantly enriched in RNA modification and ribosome biogenesis in eukaryotes (Figure 5D,E). Notably, all drought-responsive *ApWRKYs* except *ApWRKY20* were found in one of the four upregulated gene clusters: C1 (2 *WRKYs*, 448 DEGs), C2 (6 *WRKYs*, 389 DEGs), C3 (2 *WRKYs*, 294 DEGs), and C5 (5 *WRKYs*, 481 DEGs). Members of these clusters were enriched in MAPK signaling pathway, secondary metabolic biosynthetic process, cysteine and methionine metabolism, and ribosome biogenesis in eukaryotes (Figure 5D,E).

**Figure 5.**
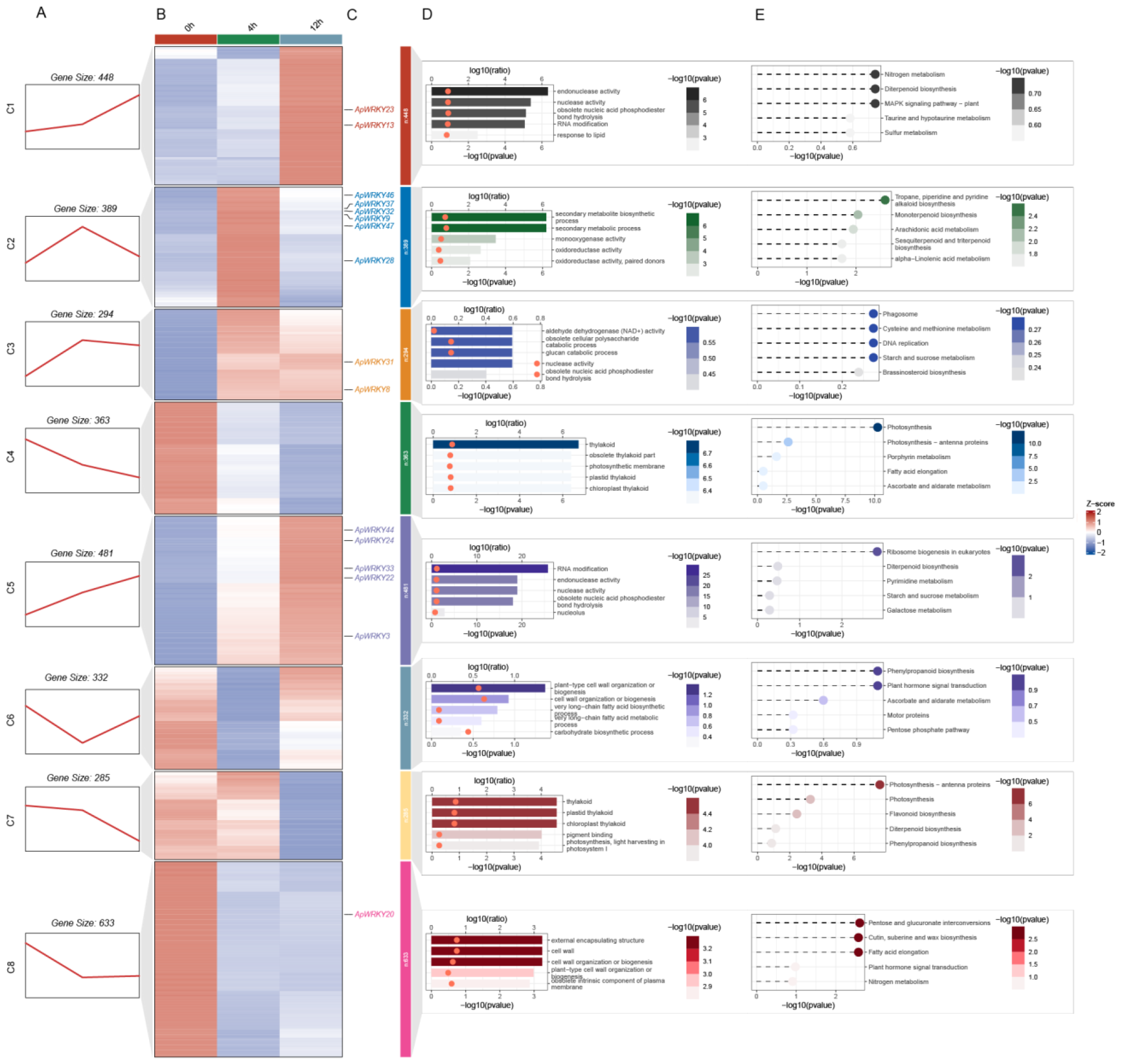
(A) Differentially expressed genes (DEGs) exhibited eight temporal expression patterns over a 12-h PEG-mediated drought treatment. (B) Clustered heatmap depicting DEG expression using TPM values with zero-mean normalization. The color scale from blue to red represents increasing TPM values. (C) *ApWRKY* genes that were differentially expressed under drought treatment. (D–E) The top five significantly enriched GO terms (D) and KEGG pathways (E) of genes in each cluster.

By contrast, *ApWRKY20* was present in C8, whose 633 members were downregulated with increasing drought duration and were significantly enriched in pentose and glucuronate interconversions, fatty acid elongation, cutin, and suberin and wax biosynthesis. Among the three smaller clusters of downregulated genes, C4 and C7 contained DEGs related to photosynthesis and thylakoids, consistent with declining photosynthetic activity under drought stress. C6 contained DEGs enriched in phenylpropanoid biosynthesis and plant hormone signal transduction whose expression first declined at 4 h and then increased again at 12 h.

Differences in their expression patterns suggest that individual *ApWRKY* genes have distinct functions and may act at different stages of the drought response. For instance, *ApWRKYs* in cluster C1 were upregulated only at the later stage of PEG-mediated drought (12 h), those in C2 were upregulated only at the earlier stage (4 h), and those in C3 and C5 were upregulated at both stages. Most upregulated *ApWRKY* genes were in either cluster C2 or C5. To better understand their potential functions, we investigated the roles of their *Arabidopsis* homologs from the phylogenetic tree in Figure 4A. AtWRKY33, the *Arabidopsis* homolog of ApWRKY28, directly and negatively regulates the cellulose synthase gene *CesA8*, resulting in improved tolerance to drought stress^37^. ApWRKY28, whose gene was upregulated at 4 h, may have a similar function. ApWRKY47 is homologous to AtWRKY50 from *Arabidopsis* and HcWRKY50 from kenaf (*Hibiscus cannabinus*), and all three proteins contain the variant WRKYGKK motif rather than the common WRKYGQK motif. A previous study demonstrated that HcWRKY50 positively regulates expression of *RD29B* and *COR47* in the ABA signaling pathway^38^, suggesting that ApWRKY47 may function similarly in ABA signaling and drought tolerance. Overexpression of the *ApWRKY22* homolog *AtWRKY30* improved heat and drought tolerance in wheat^39^. The ApWRKY33 homologs AtWRKY18/60 and the ApWRKY3 homolog AtWRKY40 form an interconnected regulatory network in *Arabidopsis* that controls the expression of genes associated with ABA, defense, and stress responses, in which AtWRKY18/60 act as transcriptional activators and *AtWRKY40* as a repressor^40^. The roles of their *Arabidopsis* homologs in drought and stress tolerance suggest that ApWRKY28, 47, 22, 33, and 3 are good candidates for future research on mechanisms of *A. pictum* responses to abiotic stress.

## Discussion

Our study presents useful genomic and transcriptomic resources for further research on mechanisms of drought resistance in *A. pictum*, a drought-tolerant medicinal plant from the Tarim Basin, and offers insights into aspects of Apocynaceae evolution. We used a combination of Illumina HiSeq, Nanopore, and Hi-C sequencing to generate a high-quality chromosome-scale genome assembly for *A. pictum*. The final assembled genome size was ∼225.3 Mb, with a scaffold N50 of 19.6 Mb and 99.55% of the genome sequences mapped to 11 pseudochromosomes. The completeness of the *A. pictum* genome assembly was confirmed by BUSCO analysis, which showed that 96.8% of the examined orthologs were completely assembled. Likewise, its high LAI score indicated that the *A. pictum* genome was of reference quality^10^.

A phylogenetic tree of selected Apocynaceae members based on high-throughput sequencing of nuclear genomic data revealed that Rauvolfioideae diverged first from the common ancestor of Rauvolfioideae, Apocynoideae, and Asclepiadoideae, consistent with previous research based on plastomes^18^. As expected, the two representative Apocynoideae species, *A. pictum* and *A. venetum*, had the closest evolutionary relationship. Through the utilization of a larger genomic dataset of the Apocynaceae family, we estimated the divergence time between *A. pictum* and *A. venetum* to be 2.2 Mya, which is earlier than previously estimated^8^. We identified 169 expanded gene families in *A. pictum*, and these genes were significantly enriched in a pathway related to environmental stress response. We also found that the tested Apocynaceae species shared only the core eudicot WGT-γ event and had not undergone additional polyploidization events. Karyotype assessments revealed that several chromosomes of *A. pictum* and *M. tenacissima* were derived completely from single AEK chromosomes, whereas *C. roseus* showed greater inter-chromosome rearrangement.

The WRKY gene family encodes a group of plant-specific transcription factors that play important roles in abiotic stress pathways^36^. We identified 50 *ApWRKYs*, established their basic classifications, and predicted the characteristics and conserved motifs of their encoded proteins. Sixteen showed changes in expression in response to PEG-mediated drought stress, all but one of which were upregulated. The abundance of ABA- and stress-related *cis*-elements in their promoters provides further evidence for the roles of ApWRKYs in regulating gene expression under abiotic stress.

In summary, the availability of a high-quality genome provides new genetic resources for future evolutionary and comparative genomics analyses of *A. pictum* and other Apocynaceae species, and transcriptomic data provide initial molecular insights into drought adaptation in this desert plant.

## Materials and methods

### Sample collection and sequencing

The individual *A. pictum* plant used for sequencing was growing in the northern region of the Tarim Basin (81.31E, 40.54N; Xinjiang, China); fresh shoots were collected and transported on dry ice to BIOYIGENE (Wuhan, China) for genome sequencing. Total genomic DNA was extracted using the QIAGEN Genomic DNA Extraction kit. After repairing DNA damage, the purified DNA fragments were subjected to end repair and A-tailing reactions at both ends of the DNA fragment. Following purification, the DNA fragments were ligated with adapters obtained from the Ligation Sequencing Kit (SQK-LSK109). Long reads were sequenced on the PromethION platform. Low-quality reads were removed after basecalling (mean_qscore_template < 7), and short reads were removed using Filtlong v0.2.1 (--min_length 1000). For short reads sequenced on the Illumina HiSeq 4000 platform, adapters and low-quality reads were removed using Trim Galore v0.6.7 (-q 25 -phred33 -length 100 -stringency 1 -paired). For Hi-C sequencing, tender shoots were fixed in a 1% formaldehyde solution to cross-link chromatin, then digested with the MboI restriction enzyme. A DNA library was constructed and sequenced on the Illumina HiSeq 4000 platform, and fastq sequencing files were trimmed with fastp^41^ v0.12.6 using default parameters. We ultimately obtained 17 Gb of Illumina short-read sequencing data, 48 Gb of Hi-C sequencing data, and 21 Gb of ONT long-read sequencing data for use in assembly of the *A. pictum* genome scaffolds (Supplementary Table S1).

Transcriptome sequencing of fresh leaf and stem tissue from the same plant was performed for the purpose of gene structural annotation. RNA extraction, library construction, and sequencing were performed by BIOYIGENE (Wuhan, China) on the MGI2000 platform. For samples from the drought experiment (see below), total RNA was extracted from leaf tissue using the QIAGEN RNAprep pure Plant Kit. Paired-end library preparation, quality control were performed, and libraries were sequenced on the MGISEQ-2000RS platform at the High-throughput Sequencing Platform of the National Key Laboratory of Crop Genetic Improvement at Huazhong Agricultural University (Wuhan, China).

### Genome survey and *de novo* assembly

We used Illumina data to estimate the genome size, heterozygosity ratio, and repetitive sequence content of *A. pictum* using k-mer distribution analysis (k = 17). We first used jellyfish v2.3.0 (-m 17 -C -s 300M) to count and compute the frequency of 17-mers and then visualized the 17-mer count histogram using GenomeScope2^42^ v2.0. We then generated a contig-level assembly of ONT reads using NextDenovo^43^ v2.5.0 (genome size = 202m, read_cutoff = 1k, correction_options = -p 15 -dbuf) and performed two rounds of polishing with Illumina short reads using NextPolish^44^ v1.4.1 with default parameters. To improve the genome contiguity and obtain a chromosome-level contig assembly, we used a combination of Juicer v2.0 (-s MboI --assembly) and 3D-DNA^45^ (two rounds of polishing with default parameters) strategies with the Hi-C sequencing data. We used Juicebox Assembly Tools^46^ to manually refine the chromosome boundaries, then executed the run-asm-pipeline-post-review.sh script in the 3D-DNA pipeline to obtain the chromosome-level assembly. We examined the pseudochromosome interactions in a Hi-C heatmap using plotHicGenome software and made manual adjustments to chromosome order and orientation.

We used multiple methods to evaluate the completeness and quality of the assembled genome, including BUSCO^9^ v5.3.2 with the eudicots_odb10 database (Supplementary Table S2) and calculation of the LAI value^10^. LTR_Finder^47^ v1.0.7 (-D 15000 -d 1000 -L 7000 -l 100 -p 20 -C -M 0.85) and LTRharvest^48^ (-minlenltr 100 -maxlenltr 7000 -mintsd 4 -maxtsd 6 -motif TGCA -motifmis 1 -similar 85 -vic 10) were used to predict LTR retrotransposons (LTR-RTs) and construct an LTR sequence library. The LAI value was then calculated using the perl program LTR_retriever^49^ with default parameters. We also used the Quality Assessment Tool^50^ v5.2.0 to align Illumina short reads to the draft genome and obtain genome statistics.

To evaluate genome base accuracy, we called SNPs and indels using the single-sample workflow of the Genome Analysis Toolkit^51^ (GATK) v4.3.0 pipeline (Supplementary Table S3). SAMtools v1.6 and bwa-mem2 v2.2.1 were used to build an index of the draft genome, and the GATK CreateSequenceDictionary script was used to create the sequence dictionary. The fastq file of Illumina short reads was converted to uBAM format with the GATK FastqToSam algorithm, then aligned to the draft genome to obtain a clean BAM file after marking Illumina adapters and duplicate sequences. A variant call format (VCF) file was generated after processing using the GATK HaplotypeCaller algorithm. SNPs and indels were identified using standard hard-filtering parameters^52^. The following parameters were used for SNPs: QD < 2.0 || MQ < 40.0 || FS > 60.0 || SOR > 3.0 || MQRankSum < -12.5 || ReadPosRankSum < -8.0. For Indels, the parameters used were: QD < 2.0 || FS > 200.0 || SOR > 10.0 || MQRankSum < -12.5 || ReadPosRankSum < -8.0.

### Repetitive element and structural annotations

We identified tandem repeats using Tandem repeats finder^53^ with the parameters “2 7 7 80 10 50 500 -f –d -m -r -h” (Supplementary Table S4). TE annotations were based on a custom repeat library that included a *de novo* repeat library generated with RepeatModeler^54^ v2.0.3, the lamiids and ancestral consensus repeats from the Repbase^55^ v20181026 database, the Dfam^56^ database, and the LTR sequence library used to calculate the LAI score. We used RepeatMasker v4.1.2 with the custom repeat library to identify repetitive elements in the genome, and we obtained the soft-masked genome with the following parameters: -nolow -no_is -engine ncbi -gff -norna -xsmall -poly (Supplementary Table S5).

To annotate non-coding RNAs (ncRNAs), we used the cmscan program (--cut_ga –rfam –nohmmonly -- fmt 2) in Infernal^57^ v1.1.4 to search for ncRNA sequences against the Rfam^58^ v14.10 database with the cmpress program. We primarily retained hits that did not overlap with any other hits or, in the case of overlapping hits, those that had a lower E-value compared with the others (Supplementary Table S6).

We performed structural annotation using the BRAKER^59^ v3.0.2 (ETPmode) automated annotation pipeline^60^. We used a custom protein database consisting of two closely related species, *Coffea arabica* (GCF_003713225.1) and *Coffea canephora* (GCA_900059795.1), for homologous prediction, and we incorporated orthologous proteins from the Viridiplantae in the OrthoDB^61^ v10 database. The custom database and 26 Gb of paired-end RNA-sequencing data were used to train GeneMark-ETP^62^ to provide supporting evidence. The soft-masked genome was then used to train AUGUSTUS^63^ for *de novo* annotation predictions using the GeneMark-ETP results (Supplementary Table S7). The final functional gene set was filtered using TEsorter^64^ v1.4.6 (-eval 1e-6), taking into consideration that these genes may have been inactivated by transposon insertion. The quality of annotations was evaluated using BUSCO^9^ in protein mode (Supplementary Table S2), and the predicted proteins were functionally annotated using the EggNOG v5.0 database^11^.

### Phylogenetic analysis

The longest transcript from ten species was used to identify orthologous gene families using OrthoFinder^65^ v2.5.5 (-M msa), with multiple sequence alignment specified through Diamond^66^ v2.1.8.162 as the method for maximum-likelihood species tree inference^67^. Divergence times for single-copy orthologs were estimated using the MCMCtree program from the PAML v4.10.6 package^68,69^. The species tree was calibrated using two fossil constraints from the TimeTree^70^ website: the estimated divergence of *O. sativa* and *A. thaliana* at 1.421–1.635 Mya and that of *A. thaliana* and *V. vinifera* at 1.098–1.244 Mya. Expansion and contraction of gene families were analyzed using CAFE5^71^ (-p -k 5) to estimate evolutionary rates under a birth–death model using maximum-likelihood estimation. Expanded gene families in *A. pictum* were subjected to GO and KEGG enrichment analyses using the ClusterProfiler^72^ v4.2.2 R package with the annotation file to predict their putative functions and pathways. Results of phylogenetic analysis were visualized using ChiPlot^73^.

### WGD analysis

Paralogous and orthologous genes within and between genomes were identified using an all-against-all homology search performed with BLASTP^74^ using an E-value cut-off of 1e−5. Syntenic blocks within and between genomes were detected using MCscanX^75^ with the criterion that at least 5 gene pairs were retained. MAFFT^76^ v7.310 multiple sequence alignment of paralogous and orthologous genes was performed with WGDI^77^ software, and the coding sequences of these genes were used to estimate non-synonymous (Ka) and synonymous (Ks) substitution rates. The NeiGojobori method implemented in the YN00 program provided by PAML was used for this purpose^69^. WGD events were detected by Gaussian fitting of the Ks distribution using WGDI software with the parameters: -bi -kp -pf -kf. Syntenic blocks between chromosomes were visualized with the bar plot function of WGDI using the parameters -km and -k.

### Identification of *WRKY* genes

The sequences of WRKY proteins from *A. thaliana* were obtained from the TAIR^29^ database. A Hidden Markov Model (HMM) search against the *A. pictum* genome was performed using the PFAM^12^ database and the CDD^78^ database to identify candidate WRKY genes. Only sequences containing the WRKY domain (PF03106) were retained as candidates. The chromosomal distribution of *ApWRKY* genes was displayed using TBtools^79^ software. From a multiple sequence alignment obtained with ClustalW using clearly classified AtWRKYs, we constructed a neighbor-joining phylogenetic tree using MEGA X^80^ with 1,000 bootstrap replicates. The ApWRKY protein sequences were submitted to MEME Suite v5.5.5 to identify up to 10 conserved motifs, and these results were integrated with the CDD search results and gene structure file using TBtools software. Protein properties of the ApWRKYs, including length, molecular weight, pI, instability index, aliphatic index, and GRAVY score, were predicted using ExPASY (Supplementary Table S10). The 2,000-bp sequences upstream of the translation initiation sites of the *ApWRKY* genes were extracted using TBtools, then submitted to the PlantCARE^81^ database to predict potential *cis*-acting regulatory elements. The results were visualized using R scripts.

### Transcriptome analysis

The healthy seedings of *A. pictum* was treated with 50ml of 30% polyethylene glycol (PEG) 6000 to simulate drought stress conditions throughout their 4-week growth period. Drought stress was divided into two stages: early(4 hours) and late(12 hours), and each treatment replicated for three times. Leaf tissues collected from *A. pictum* under 0, 4, and 12 h drought stress were used to prepare Illumina sequencing libraries as described above. Adapters and low-quality reads were removed from the resulting sequencing data using fastp^41^ v0.12.6 with default parameters. The clean reads were mapped to the *A. pictum* genome, and raw count values were calculated using STAR^82^ v2.5.2 (--quantMode GeneCounts). Differentially expressed genes were defined as those exhibiting at least one-fold change in expression compared to the 0-h time point and were identified using the DESeq2 R package, with an FDR-adjusted p-value <0.05. To investigate the expression patterns of DEGs at two time points, we transformed the raw count data into transcripts per kilobase of exon model per million mapped reads (TPM). Cluster analysis of gene expression patterns was performed with the Mfuzz^83^ v2.54.0 R package; the results were used to construct an expression heatmap and perform KEGG/GO enrichment analysis using the ClusterGvis R package^84^.

## Acknowledgements

This work was supposed by the Construction of the Innovation, Utilization, and Demonstration Promotion of Stress-resistant Cotton Varieties in South Xinjiang program (2022DB012) and the Molecular Design Breeding System for Stress Resistance and High Yield Production program (2023ZD04040-4-2). We received support for computational work from the High-throughput Sequencing Platform of the National Key Laboratory of Crop Genetic Improvement in Huazhong Agricultural University and the Bioinformatics Computing Platform in Tarim University. We were also grateful to the members of Maojun Wang’s research group in Huazhong Agricultural University for their technical advice during the assembly and annotation steps.

## Contributions

Y.Q.W. conceived and designed the project. B.W.B. collected the genomic materials. W.L.X. assembled the genome and performed gene annotation, evolutionary analyses, WRKY family analyses, and conducted drought treatment experiment and transcriptomic analyses. W.L.X., B.W.B. and Y.Q.W. wrote the manuscript. All authors read and approved the final version of the manuscript.

## Data availability

All raw sequence reads produced in this study, including Illumina, Nanopore, and Hi-C interaction reads, and RNA-seq data from drought treatments, have been deposited at the NCBI Sequence Read Archive (SRA) under BioProject PRJNA1069313. Pertinent datasets, comprising the final assembled genome and its annotations, have been deposited at Figshare with the DOI 10.6084/m9.figshare.25060931.

## References

1 Sun, J., Zhang, L., Deng, C. & Zhu, R. Evidence for enhanced aridity in the Tarim Basin of China since 5.3 Ma. Quaternary Science Reviews 27, 1012–1023 (2008).

2 Jiang, N., Zhang, Q., Zhang, S., Zhao, X. & Cheng, H. Spatial and temporal evolutions of vegetation coverage in the Tarim River Basin and their responses to phenology. Catena 217, 106489 (2022).

3 Yang, J., Zhang, L., Jiang, L., Zhan, Y. G. & Fan, G. Z. Quercetin alleviates seed germination and growth inhibition in Apocynum venetum and Apocynum pictum under mannitol-induced osmotic stress. Plant Physiol Biochem 159, 268-276, doi:10.1016/j.plaphy.2020.12.025 (2021).

4 Rouzi, A., Halik, Ü., Thevs, N., Welp, M. & Aishan, T. Water efficient alternative crops for sustainable agriculture along the Tarim basin: a comparison of the economic potentials of Apocynum pictum, Chinese red date and cotton in Xinjiang, China. Sustainability 10, 35 (2017).

5 Ma, M., Hong, C.-L., An, S.-Q. & Li, B. Seasonal, spatial, and interspecific variation in quercetin in Apocynum venetum and Poacynum hendersonii, Chinese traditional herbal teas. Journal of agricultural and food chemistry 51, 2390–2393 (2003).

6 4 Xie, W., Zhang, X., Wang, T. & Hu, J. Botany, traditional uses, phytochemistry and pharmacology of Apocynum venetum L.(Luobuma): A review. Journal of Ethnopharmacology 141, 1–8 (2012).

7 Zheng, C. et al. The complete chloroplast genome and phylogenetic relationship of Apocynum pictum (Apocynaceae), a Central Asian shrub and second-class national protected species of western China. Gene 830, 146517 (2022).

8 Gao, G. et al. Comparative genome and metabolome analyses uncover the evolution and flavonoid biosynthesis between Apocynum venetum and Apocynum hendersonii. Iscience 26 (2023).

9 Simao, F. A., Waterhouse, R. M., Ioannidis, P., Kriventseva, E. V. & Zdobnov, E. M. BUSCO: assessing genome assembly and annotation completeness with single-copy orthologs. Bioinformatics 31, 3210–3212, doi:10.1093/bioinformatics/btv351 (2015).

10 Ou, S., Chen, J. & Jiang, N. Assessing genome assembly quality using the LTR Assembly Index (LAI). Nucleic acids research 46, e126–e126 (2018).

11 Huerta-Cepas, J. et al. eggNOG 5.0: a hierarchical, functionally and phylogenetically annotated orthology resource based on 5090 organisms and 2502 viruses. Nucleic acids research 47, D309–D314 (2019).

12 Mistry, J. et al. Pfam: The protein families database in 2021. Nucleic acids research 49, D412–D419 (2021).

13 Xu, Z. et al. A near-complete genome assembly of Catharanthus roseus and insights into its vinblastine biosynthesis and high susceptibility to the Huanglongbing pathogen. Plant Communications 4 (2023).

14 Cuello, C. et al. Genome assembly of the medicinal plant Voacanga thouarsii. Genome Biology and Evolution 14, evac158 (2022).

15 Zhou, Y. et al. Marsdenia tenacissima genome reveals calcium adaptation and tenacissoside biosynthesis. The Plant Journal 113, 1146–1159 (2023).

16 Weitemier, K. et al. A draft genome and transcriptome of common milkweed (Asclepias syriaca) as resources for evolutionary, ecological, and molecular studies in milkweeds and Apocynaceae. PeerJ 7, e7649 (2019).

17 Hoopes, G. M. et al. Genome assembly and annotation of the medicinal plant Calotropis gigantea, a producer of anticancer and antimalarial cardenolides. G3: Genes, Genomes, Genetics 8, 385–391 (2018).

18 Fishbein, M. et al. Evolution on the backbone: Apocynaceae phylogenomics and new perspectives on growth forms, flowers, and fruits. American Journal of Botany 105, 495–513 (2018).

19 Del Rio, C. et al. Asclepiadospermum gen. nov., the earliest fossil record of Asclepiadoideae (Apocynaceae) from the early Eocene of central Qinghai-Tibetan Plateau, and its biogeographic implications. American Journal of Botany 107, 126–138 (2020).

20 Ribeiro, p. L., Rapini, A., Damascena, L. S. & van den Berg, C. Plant diversification in the Espinhaço Range: insights from the biogeography of Minaria (Apocynaceae). Taxon 63, 1253–1264 (2014).

21 Bitencourt, C. et al. Evolution of dispersal, habit, and pollination in Africa pushed Apocynaceae diversification after the Eocene-Oligocene climate transition. Frontiers in Ecology and Evolution 9, 719741 (2021).

22 Liu, J.-X. & Howell, S. H. Endoplasmic reticulum protein quality control and its relationship to environmental stress responses in plants. The Plant Cell 22, 2930–2942 (2010).

23 Kachroo, A. & Kachroo, p. Fatty acid–derived signals in plant defense. Annual review of phytopathology 47, 153–176 (2009).

24 4 Gao, G. et al. UPLC-ESI-MS/MS based characterization of active flavonoids from apocynum spp. and anti-bacteria assay. Antioxidants 10, 1901 (2021).

25 Wang, Z. et al. A high-quality Buxus austro-yunnanensis (Buxales) genome provides new insights into karyotype evolution in early eudicots. BMC biology 20, 1–17 (2022).

26 Salse, J. Ancestors of modern plant crops. Current Opinion in Plant Biology 30, 134–142 (2016).

27 The grapevine genome sequence suggests ancestral hexaploidization in major angiosperm phyla. nature 449, 463–467 (2007).

28 Zhang, Y. & Wang, L. The WRKY transcription factor superfamily: its origin in eukaryotes and expansion in plants. BMC evolutionary biology 5, 1–12 (2005).

29 Berardini, T. Z. et al. The Arabidopsis information resource: making and mining the “gold standard” annotated reference plant genome. genesis 53, 474–485 (2015).

30 Mohan, R. & Venugopal, S. Computational structural and functional analysis of hypothetical proteins of Staphylococcus aureus. Bioinformation 8, 722 (2012).

31 Eulgem, T., Rushton, p. J., Robatzek, S. &Somssich, I. E. The WRKY superfamily of plant transcription factors. Trends in plant science 5, 199–206 (2000).

32 Xie, Z. et al. Annotations and functional analyses of the rice WRKY gene superfamily reveal positive and negative regulators of abscisic acid signaling in aleurone cells. Plant physiology 137, 176–189 (2005).

33 Ross, C. A., Liu, Y. & Shen, Q. J. The WRKY gene family in rice (Oryza sativa). Journal of Integrative Plant Biology 49, 827–842 (2007).

34 Wan, Y. et al. Identification of the WRKY gene family and functional analysis of two genes in Caragana intermedia. BMC plant biology 18, 1–16 (2018).

35 Liu, J., Li, G., Wang, R., Wang, G. & Wan, Y. Genome-Wide Analysis of WRKY Transcription Factors Involved in Abiotic Stress and ABA Response in Caragana korshinskii. International Journal of Molecular Sciences 24, 9519 (2023).

36 Li, W., Pang, S., Lu, Z. & Jin, B. Function and mechanism of WRKY transcription factors in abiotic stress responses of plants. Plants 9, 1515 (2020).

37 Wang, X., Du, B., Liu, M., Sun, N. & Qi, X. Arabidopsis transcription factor WRKY33 is involved in drought by directly regulating the expression of CesA8. (2013).

38 Niu, X. et al. Ectopic Expression of Kenaf (Hibiscus cannabinus L.) HcWRKY50 Improves Plants’ Tolerance to Drought Stress and Regulates ABA Signaling in Arabidopsis. Agronomy 12, 1176 (2022).

39 El-Esawi, M. A., Al-Ghamdi, A. A., Ali, H. M. & Ahmad, M. Overexpression of AtWRKY30 transcription factor enhances heat and drought stress tolerance in wheat (Triticum aestivum L.). Genes 10, 163 (2019).

40 Chen, H. et al. Roles of Arabidopsis WRKY18, WRKY40 and WRKY60 transcription factors in plant responses to abscisic acid and abiotic stress. BMC plant biology 10, 1–15 (2010).

41 Chen, S., Zhou, Y., Chen, Y. & Gu, J. fastp: an ultra-fast all-in-one FASTQ preprocessor. Bioinformatics 34, i884–i890, doi:10.1093/bioinformatics/bty560 (2018).

42 Ranallo-Benavidez, T. R., Jaron, K. S. & Schatz, M. C. GenomeScope 2.0 and Smudgeplot for reference-free profiling of polyploid genomes. Nat Commun 11, 1432, doi:10.1038/s41467-020-14998-3 (2020).

43 Hu, J. et al. An efficient error correction and accurate assembly tool for noisy long reads. bioRxiv, 2023.2003.2009.531669, doi:10.1101/2023.03.09.531669 (2023).

44 Hu, J., Fan, J., Sun, Z. & Liu, S. NextPolish: a fast and efficient genome polishing tool for long-read assembly. Bioinformatics 36, 2253–2255 (2020).

45 Dudchenko, O. et al. De novo assembly of the Aedes aegypti genome using Hi-C yields chromosome-length scaffolds. Science 356, 92–95, doi:10.1126/science.aal3327 (2017).

46 Durand, N. C. et al. Juicebox provides a visualization system for Hi-C contact maps with unlimited zoom. Cell systems 3, 99–101 (2016).

47 Xu, Z. & Wang, H. LTR_FINDER: an efficient tool for the prediction of full-length LTR retrotransposons. Nucleic acids research 35, W265–W268 (2007).

48 Ellinghaus, D., Kurtz, S. & Willhoeft, U. LTRharvest, an efficient and flexible software for de novo detection of LTR retrotransposons. BMC bioinformatics 9, 1–14 (2008).

49 Ou, S. & Jiang, N. LTR_retriever: a highly accurate and sensitive program for identification of long terminal repeat retrotransposons. Plant physiology 176, 1410–1422 (2018).

50 Gurevich, A., Saveliev, V., Vyahhi, N. & Tesler, G. QUAST: quality assessment tool for genome assemblies. Bioinformatics 29, 1072–1075 (2013).

51 McKenna, A. et al. The Genome Analysis Toolkit: a MapReduce framework for analyzing next-generation DNA sequencing data. Genome research 20, 1297–1303 (2010).

52 O’Connor, B. D. & van der Auwera, G. Genomics in the Cloud: Using Docker, GATK, and WDL in Terra. (O’Reilly Media, Incorporated, 2020).

53 Benson, G. Tandem repeats finder: a program to analyze DNA sequences. Nucleic acids research 27, 573–580 (1999).

54 Flynn, J. M. et al. RepeatModeler2 for automated genomic discovery of transposable element families. Proceedings of the National Academy of Sciences 117, 9451–9457 (2020).

55 Jurka, J. et al. Repbase Update, a database of eukaryotic repetitive elements. Cytogenetic and genome research 110, 462–467 (2005).

56 Storer, J., Hubley, R., Rosen, J., Wheeler, T. J. & Smit, A. F. The Dfam community resource of transposable element families, sequence models, and genome annotations. Mobile DNA 12, 1–14 (2021).

57 Nawrocki, E. p. & Eddy, S. R. Infernal 1.1: 100-fold faster RNA homology searches. Bioinformatics 29, 2933–2935, doi:10.1093/bioinformatics/btt509 (2013).

58 Kalvari, I. et al. Rfam 14: expanded coverage of metagenomic, viral and microRNA families. Nucleic Acids Research 49, D192–D200, doi:10.1093/nar/gkaa1047 (2020).

59 Gabriel, L. et al. BRAKER3: Fully automated genome annotation using RNA-Seq and protein evidence with GeneMark-ETP, AUGUSTUS and TSEBRA. bioRxiv, doi:10.1101/2023.06.10.544449 (2023).

60 Hoff, K. J., Lomsadze, A., Borodovsky, M. & Stanke, M. Whole-genome annotation with BRAKER. Gene prediction: methods and protocols, 65–95 (2019).

61 Kuznetsov, D. et al. OrthoDB v11: annotation of orthologs in the widest sampling of organismal diversity. Nucleic Acids Res 51, D445–D451, doi:10.1093/nar/gkac998 (2023).

62 Bruna, T., Lomsadze, A. & Borodovsky, M. GeneMark-EP+: eukaryotic gene prediction with self-training in the space of genes and proteins. NAR Genom Bioinform 2, lqaa026, doi:10.1093/nargab/lqaa026 (2020).

63 Stanke, M. et al. AUGUSTUS: ab initio prediction of alternative transcripts. Nucleic acids research 34, W435–W439 (2006).

64 Zhang, R.-G. et al. TEsorter: an accurate and fast method to classify LTR-retrotransposons in plant genomes. Horticulture Research 9, uhac017 (2022).

65 Emms, D. M. & Kelly, S. OrthoFinder: phylogenetic orthology inference for comparative genomics. Genome biology 20, 1–14 (2019).

66 Buchfink, B., Reuter, K. & Drost, H.-G. Sensitive protein alignments at tree-of-life scale using DIAMOND. Nature methods 18, 366–368 (2021).

67 Emms, D. & Kelly, S. STAG: species tree inference from all genes. BioRxiv, 267914 (2018).

68 Reis, M. d. & Yang, Z. Approximate likelihood calculation on a phylogeny for Bayesian estimation of divergence times. Molecular biology and evolution 28, 2161–2172 (2011).

69 Yang, Z. PAML 4: phylogenetic analysis by maximum likelihood. Molecular biology and evolution 24, 1586–1591 (2007).

70 Kumar, S., Stecher, G., Suleski, M. & Hedges, S. B. TimeTree: A Resource for Timelines, Timetrees, and Divergence Times. Molecular Biology and Evolution 34, 1812-1819, doi:10.1093/molbev/msx116 (2017).

71 Mendes, F. K., Vanderpool, D., Fulton, B. & Hahn, M. W. CAFE 5 models variation in evolutionary rates among gene families. Bioinformatics 36, 5516–5518 (2020).

72 Wu, T. et al. clusterProfiler 4.0: A universal enrichment tool for interpreting omics data. The innovation 2 (2021).

73 Xie, J. et al. Tree Visualization By One Table (tvBOT): a web application for visualizing, modifying and annotating phylogenetic trees. Nucleic Acids Research 51, W587–W592, doi:10.1093/nar/gkad359 (2023).

74 Camacho, C. et al. BLAST+: architecture and applications. BMC bioinformatics 10, 1–9 (2009).

75 Wang, Y. et al. MCScanX: a toolkit for detection and evolutionary analysis of gene synteny and collinearity. Nucleic acids research 40, e49–e49 (2012).

76 Katoh, K. & Standley, D. M. MAFFT multiple sequence alignment software version 7: improvements in performance and usability. Molecular biology and evolution 30, 772–780 (2013).

77 Sun, p. et al. WGDI: A user-friendly toolkit for evolutionary analyses of whole-genome duplications and ancestral karyotypes. Molecular plant 15, 1841–1851 (2022).

78 Wang, J. et al. The conserved domain database in 2023. Nucleic Acids Res 51, D384–D388, doi:10.1093/nar/gkac1096 (2023).

79 Chen, C. et al. TBtools: an integrative toolkit developed for interactive analyses of big biological data. Molecular plant 13, 1194–1202 (2020).

80 Kumar, S., Stecher, G., Li, M., Knyaz, C. & Tamura, K. MEGA X: Molecular Evolutionary Genetics Analysis across Computing Platforms. Mol Biol Evol 35, 1547-1549, doi:10.1093/molbev/msy096 (2018).

81 Lescot, M. et al. PlantCARE, a database of plant cis-acting regulatory elements and a portal to tools for in silico analysis of promoter sequences. Nucleic acids research 30, 325–327 (2002).

82 Dobin, A. et al. STAR: ultrafast universal RNA-seq aligner. Bioinformatics 29, 15-21, doi:10.1093/bioinformatics/bts635 (2013).

83 Kumar, L. & Futschik, M. E. Mfuzz: a software package for soft clustering of microarray data. Bioinformation 2, 5 (2007).

84 Zhang, J. ClusterGVis: One-step to Cluster and Visualize Gene Expression Matrix., <https://github.com/junjunlab/ClusterGVis> (2022).

